# Cardiac cycle affects the asymmetric value updating in instrumental reward learning

**DOI:** 10.1101/2022.03.03.482830

**Authors:** Kenta Kimura, Noriaki Kanayama, Asako Toyama, Kentaro Katahira

## Abstract

This study aimed to investigate whether instrumental reward learning is affected by the cardiac cycle. To this end, we examined the effects of the cardiac cycle (systole or diastole) on the computational processes underlying the participants’ choices in the instrumental learning task. In the instrumental learning task, participants were required to select one of two discriminative stimuli (neutral visual stimuli) and immediately receive reward/punishment feedback depending on the probability assigned to the chosen stimuli. To manipulate the cardiac cycle, the presentation of discriminative stimuli was timed to coincide with either cardiac systole or diastole. We fitted the participants’ choices in the task with reinforcement learning (RL) models and estimated parameters involving instrumental learning (i.e., learning rate and inverse temperature) separately in the systole and diastole trials. Model-based analysis revealed that the learning rate for positive prediction errors was higher than that for negative prediction errors in the systole trials; however, learning rates did not differ between positive and negative prediction errors in the diastole trials. These results demonstrate that the natural fluctuation of cardiac afferent signals can affect asymmetric value updating in instrumental reward learning.

## Introduction

It is widely accepted that not only does the brain regulate the internal physiological state of the body, but information concerning the internal physiological state of the body is also transmitted to the brain. This bi-directional signal processing between the brain and the internal physiological state of the body is called “Interoception” (e.g., Chen et al., 2021). Interoceptive signals are originated from all major biological systems, including the cardiovascular, gastrointestinal, immune, and autonomic systems (for a comprehensive review, see Khalsa et al., 2018). Recent theoretical and empirical research has provided converging evidence that interoception plays an essential role in energy regulation, subjective sense of self, and affective experience (for a review, see Quingley et al., 2021).

In previous studies on interoception, afferent signals from heart activity (i.e., cardiac afferent signals) have been the target of a growing body of research. The strength and timing of arterial pressure at each heartbeat are encoded by the phasic discharge of arterial baroreceptors during cardiac systole and the contraction of the heart, which is transmitted to the brainstem and used for the baroreflex control of blood pressure. Importantly, the cardiac afferent signals from the arterial baroreceptors are conveyed to areas of the brain associated with the processing of cognitive and affective information (for a review, see Garfinkel & Critchley, 2016). Consistent with this, previous studies have found that natural fluctuations in cardiac afferent signals influence the processing of several types of external stimuli. Specifically, recent studies have accumulated evidence indicating that the processing of external stimuli can be facilitated during cardiac systole, especially when the stimuli are associated with motivational or affective significance. Garfinkel et al. (2014) found that the detection of threat-related stimuli (i.e., a fearful face) in the attentional blink task was enhanced when the stimuli were presented during cardiac systole compared to cardiac diastole. Similarly, other studies have shown an enhancement of attentional capture for threat-related stimuli presented during cardiac systole in the attentional engagement task (Azevedo et al., 2018). In addition, recent studies have reported that processing positively valenced stimuli (i.e., monetary rewards and happy faces) can be facilitated during cardiac systole (Kimura, 2019; Leganes-Fonteneau et al., 2021). Therefore, previous results have suggested that the natural fluctuation of cardiac afferent signals causes moment-to-moment fluctuations in the processing of stimuli associated with motivational/affective significance.

Although previous studies have demonstrated that the cardiac cycle influences affective processing, its effect of the cardiac cycle on learning remains unclear. It is widely accepted that attention to and processing of conditioned stimuli in Pavlovian learning or discriminative stimuli in instrumental learning play a prominent role in a variety of learning contexts (e.g., Anderson et al., 2011; Mackintosh, 1975). Therefore, considering that the cardiac cycle modulates the processing of stimuli associated with motivational/affective significance, it is reasonable to expect that the cardiac cycle affects learning. Only one study, that is, Waselius et al. (2018), examined this issue. In their study, human participants and rabbits were subjected to trace eyeblink conditioning, in which the tone was a conditioned stimulus and an air puff toward the eye was an unconditioned stimulus. The onset of delivery of the conditioned stimulus coincided with either cardiac systole or diastole. The authors reported that the cardiac cycle modulated neural responses to the conditioned stimulus in both humans and rabbits and influenced Pavlovian learning in rabbits. Their results suggest that the cardiac cycle affects Pavlovian learning by modulating the processing of conditioned stimuli. However, no study has examined the effect of cardiac cycle on instrumental learning. Mackintosh (1975) proposed that attention to discriminative stimuli influences changes in associative strength during instrumental learning. From this perspective, it is possible that the cardiac cycle can modulate the processing of discriminative stimuli, and, hence, can affect instrumental reward learning.

To examine this possibility, this study aimed to investigate whether the cardiac cycle affects instrumental reward-learning. For this purpose, we used a model-based approach and examined the effects of the cardiac cycle on the computational processes underlying the participants’ choices in the instrumental learning task. We employed RL models that have been successfully used to capture a broad range of value-based learning at the level of both behavior and neural signals (for a review, see O’Doherty et al., 2007). In the instrumental learning task, participants could choose one of two neutral visual stimuli and immediately receive reward or punishment feedback, depending on the probability assigned to the chosen stimuli. According to a previous study investigating the effect of the cardiac cycle on learning (Waselius et al., 2018), we manipulated the onset of the presentation of discriminative stimuli (i.e., two neutral visual stimuli) such that they coincided with either cardiac systole or diastole across trials (i.e., systole and diastole trials). We fitted the participants’ choices in the task with RL models and estimated the parameters involving instrumental learning (i.e., learning rate and inverse temperature) separately in the systole and diastole trials. The difference in the estimated parameters between the systole and diastole trials was then examined. If the cardiac cycle affects instrumental reward learning, the estimated parameters would differ in the systole and diastole trials. In contrast, if the cardiac cycle did not affect instrumental reward learning, the estimated parameters would not differ between the trials.

## Methods

### Participants

Overall, 45 adults participated in our experiment (13 women, 32 men, age range = 20–42 years, mean = 24.0 years). All participants were right-handed, had normal or corrected-to-normal vision and no history of neurological, cardiovascular, or mental disorders. The experimental procedures were approved by the Safety and Ethics Committee of the National Institute of Advanced Industrial Science and Technology (AIST). All participants understood the details of the experiment before their participation, and written informed consent was obtained from each participant before the experiment. This research was conducted in accordance with ethical regulations. One participant was excluded because of technical problems. Eight participants were excluded because their performance was not significantly different from chance (binomial test, *p* > .05). Thus, the final dataset comprised 36 participants (8 females, 28 males, age range = 20–42 years, mean = 24.0 years). The final sample size was almost equivalent to that of previous studies examining the effect of the cardiac cycle on cognitive and affective processing (e.g., Azevedo et al., 2017; Kimura, 2019; Kimura et al., 2022; Waselius et al., 2018).

### Electrocardiograms (ECG) recording

ECG was recorded using an MP150 Biopac System (ECG100C). The ECG was recorded with Ag/AgCl electrodes placed on the right collarbone and the left rib. The sampling rate was 2,000 Hz, and a hardware bandpass filter between .3 and 1,000 Hz was applied. The signal was recorded using the AcqKnowledge software (Biopac Systems).

To synchronize the onset of the presentation of the discriminative stimuli, heartbeats were detected online using a threshold-based R-peak detection method in AcqKnowledge software. Using the timing of each heartbeat, the onset of presentation of the discriminative stimuli was set to coincide with the systolic (~300 ms after the R-peak) or diastolic (~550 ms after the R-peak) phases of the cardiac cycle (Azevedo et al., 2017, 2018; Garfinkel et al., 2014; Gray et al., 2009).

### Setup and experimental task

In the instrumental learning task, participants selected one of two neutral visual stimuli (at least for our participants) repeatedly for 360 trials, with minutes of break after every 40 trials. In each of the 360 trials, participants received reward feedback (10 Japanese yen) or punishment feedback (−10 Japanese yen) depending on the probability assigned to the chosen option. The Presentation software (Neurobehavioral Systems installed on a Lenovo, ThinkPad W540 computer laptop) presented the stimuli and recorded participants’ responses. All the visual stimuli were presented on a 22-inch LCD monitor (Dell E2210).

Figure 1 shows the flowchart of each trial. Each trial began with a fixation display with a black cross presented at the center of a gray background. The duration of the fixation display was controlled, trial by trial, to adjust the onset of presentation of the discriminative stimuli, such that the duration of the fixation display was 1,000 ms on average and remained the same in all cardiac cycle trials. The fixation display was followed by the display of two discriminative stimuli (i.e., stimuli A and B) presented on either side of the fixation cross. The positions of the two stimuli (left and right) were randomized. Participants were required to choose one stimulus by pressing the left button (i.e., “Z” button on a keyboard) with the index finger or the right button (“X” button on a keyboard) with the middle finger, using their dominant hand. No time limits for choices were set, and the two stimuli remained until the participant pressed the button. After the participant’s response, the chosen stimulus was highlighted by a thickening of the black outline. After 1,000 ms, reward/punishment feedback was displayed, in which the fixation cross was replaced by a colored circle (red or blue). The mapping of red/blue and reward/punishment was counterbalanced across the participants. The inter-trial interval was 1,500–2,500 msec.

**Figure 1.**
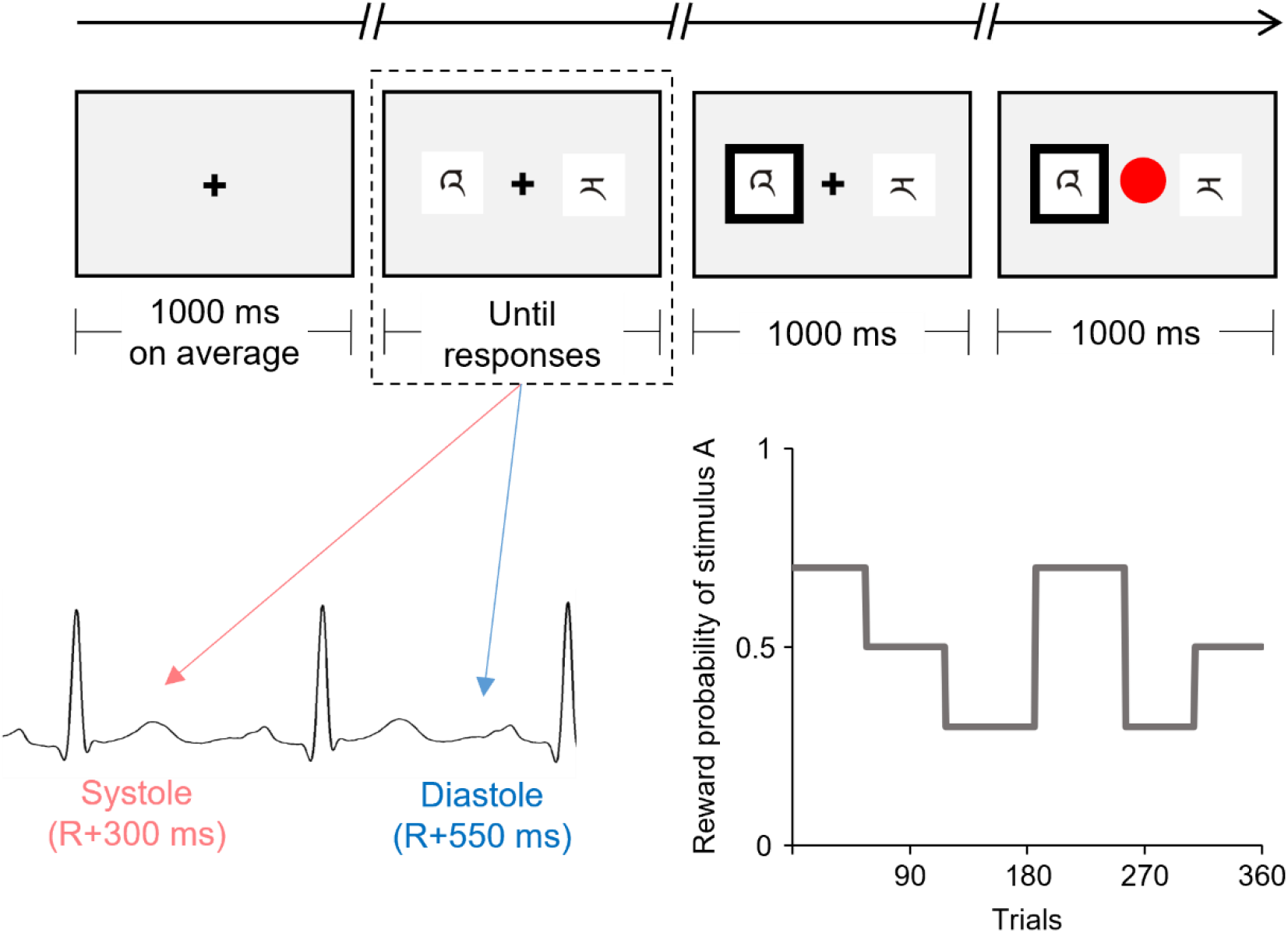
Schematic illustration of the flow of one trial of the instrumental learning task. The presentation of the discriminative stimuli (indicated by the dashed line) was experimentally manipulated to coincide with either the cardiac systole or diastole. This figure shows the reward probability for one of the two discriminative stimuli (stimulus A) which was changed among 6 blocks according to the pre-determined schedule.

The onset of the display of discriminative stimuli was synchronized to coincide with either the participant's cardiac systole or diastole. Half of the trials (180 trials) were synchronized to coincide with the cardiac systole ( systole trials), whereas the other half were synchronized to coincide with cardiac diastole (diastole trials).

The reward probability for each stimulus was unknown to the participants and changed among the six blocks according to the predetermined schedule (see Figure 1). The number of trials in each block ranged from 52–69. The reward probabilities for stimuli A vs. B were 70% vs. 30%, 50% vs. 50%, or 30% vs. 70% for each block. Therefore, the participants were required to continuously monitor the contingency between choice and reward feedback over the course of the task to maximize their reward earnings. To directly compare the time courses for the choices and the effect of the cardiac cycle, all participants were confronted with the same reward probabilities. The participants were instructed that the reward probabilities could change during the task, but received no information as to how often such a change might occur.

### Procedure

Upon arrival, the participants were informed about the experiment, and asked to provide informed consent. After their height and weight were measured, the participants were seated comfortably in front of the display, and the electrodes for the ECG were attached. The participants were then asked to relax for five minutes to familiarize themselves with the laboratory environment and the electrodes. After the participants received instructions regarding the instrumental learning task, they were given a practice block of 10 trials to familiarize themselves with the task. Then, the participants received an instruction about the monetary reward: they were told that (a) they would earn 10 Japanese yen for each reward feedback and lose 10 Japanese yen for punishment feedback and (b) they would receive a cumulative reward for the entire task. After the instruction was given, the participants performed the task, which consisted of 360 trials, with minutes of break after every 40 trials. At the end of the experiment, the participants received a predefined participation fee of 5,000 Japanese yen with a task-related bonus.

### Behavioral data analysis

Reaction time was measured as the latency in milliseconds between the onset of presentation of the discriminative stimuli and when the button was pressed in each trial. The proportion of choices for Stimulus A was calculated for each reward probability.

### Model-based analysis

To examine the effect of the cardiac cycle on the parameter estimates derived from the computational RL model, the following procedure was adopted: first, to capture the participants’ choice behavior in the present task, we constructed a model set in which the effect of the cardiac cycle was not considered, fitted them to the participants’ choice data, and determined the best-fitting model. Second, using the best-fitting model, different parameters in the systole and diastole trials were estimated. We focus on two learning rates (*α*^+^ and *α*^−^, see below) and inverse temperature *β*, and examine whether these parameter estimates are different between the systole and diastole trials.

#### Q-learning models

We constructed computational models and fitted them to the participants’ choice data for the instrumental learning task. We employed a conventional reinforcement learning model termed the Q-learning model (Sutton & Barto, 1998). In the standard Q-learning model, the action value for the chosen option (e.g., stimulus A) in trial *t*, denoted by Q_*c*_ (*t*), is updated based on the following equation:

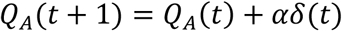

Where α is the learning rate that determines the degree of the update. *δ*(*t*) represents the prediction error which is calculated as:

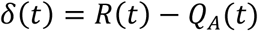

Here, *R*(*t*) is the outcome obtained by choosing stimulus A in trial *t*. The prediction error represents the difference between the expected and actual outcomes. Therefore, the action values for each option increase when the obtained outcome is better than expected, whereas they decrease when the obtained outcome is worse than expected. The probability of choosing stimulus A is given by the set of Q values according to the following softmax rule:

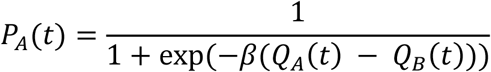

Here, *Q*_*B*_(*t*) is the action value for choosing stimulus B at trial *t*. *β* is the inverse temperature that determines the degree of stochasticity in the decision-making process.

Since previous studies have demonstrated that the effect of the cardiac cycle on affective processing could depend on stimulus valence (e.g., Azvedo et al., 2018; Garfinkel et al., 2014; Kimura, 2019; Leganes-Fonteneau et al., 2021), we used a modified version of the Q-learning model in which learning from positive and negative prediction errors is determined by different learning rates, according to previous studies (e.g., Lefebre et al., 2017). The modified version of the Q-learning model, referred to as the Q-A model, allows the learning rates to differ depending on the sign of the prediction error, as follows:

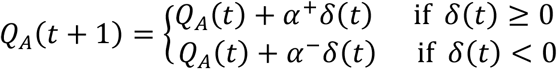

The learning rate *α*^+^ scales the extent to which the model updates the action value from one trial to the next when the prediction error is positive, whereas *α*^−^ is the same when the prediction error is negative.

We also considered three variants of the Q-A model (Q-AF, Q-AC, and Q-AFC) as candidate models. Table 1 presents the model details. The Q-AF model instantiates value-forgetting, where the action value for the unchosen option is updated by the forgetting parameter *α*_*F*_ (e.g., Katahira et al., 2017; Toyama et al., 2017). The Q-AC model includes the perseverance factor, which introduces the effect of a past choice to the choice probability (e.g., Sugawara & Katahira, 2021). The Q-AFC model combines the Q-AF and Q-AC models. We also included the null model, which is a biased random choice model that produces the same probability of two options being chosen with biases in the participants’ choices. Five models were used to fit the choice data.

**Table 1.** Information concerning the five models compared on the basis of their fit to the choice data from 36 participants.

| Model name | Description | # of free parameters | LML |
| --- | --- | --- | --- |
| Q-A | The standard Q-learning model with asymmetric learning rates ($\alpha^+$ and $\alpha^-$) for positive and negative reward prediction errors | 3 | -211.5<br>(6.54) |
| Q-AF | The Q-A model with updating unchosen action values using forgetting parameter | 4 | -205.6<br>(7.34) |
| Q-AC | The Q-A model with the computational process of choice history using decay rate ($\tau$) and perseverance parameter ($\phi$) | 5 | -206.5<br>(7.28) |
| Q-AFC | The hybrid of the Q-AF and Q-AC models | 6 | -207.3<br>(7.37) |
| null mode | The biased random choice model producing the same probability of two options being chosen with biases of the participants' choices | 1 | -248.5<br>(0.75) |

This list of models shows the mean values and standard errors across participants regarding the log marginal likelihood (LML) for each model.

#### Parameter estimation and model comparison

We used the R function “solnp” in the Rsolnp package to estimate the parameters of each model with the maximum a posteriori (MAP) estimation and calculated the log marginal likelihood of each model using Laplace approximation (Kass & Raftery, 1995). Marginal likelihood penalizes a complex model with additional parameters in the marginalization process. As the marginal likelihood is proportional to the posterior probability of the model, a higher marginal likelihood indicates a better model. Notably, this situation is true only if all models have an equal prior probability (i.e., all models are equally likely before the data are provided). This method incorporates prior distributions of the parameters and avoids extreme values in parameter estimates. Prior distributions and constraints were set according to Niv et al. (2012) and Sugawara and Katahira (2021), since these previous studies used RL models similar to this study and successfully captured the participants’ choice behavior in reward learning tasks. As prior distributions, we used a beta distribution with hyperparameters (*a* = 2, *b* = 2) for all learning rates, forgetting parameter, and decay rate, a gamma distribution (shape = 2, scale = 3) for the inverse temperature *β*, and a normal distribution (*μ* = 0, *σ*^2^ = 5) for the perseverance parameter *ϕ*. All learning rates and forgetting parameters were constrained to the range of 0 ≤ α ≤ 1. The inverse temperature was constrained to *β* ≥ 0. In the perseverance model, the decay rate was constrained to the range of 0 ≤ *τ* ≤ 1, and the perseverance parameter was constrained to the range of − 10 ≤ *ϕ* ≤ 10.

The model parameters (*α*^+^, *α*^−^, and *β*) were compared between the systole and diastole trials. Learning rates were subjected to a two-way repeated-measures ANOVA, with two trial types (systole and diastole) × two learning rate types (+ and −). The inverse temperatures for the systole and diastole trials were analyzed using *t-tests*. Effect sizes were calculated using Cohen’s *d*. An alpha level of .05 was used for all statistical analyses. The difference in learning rate asymmetry (*α*^+^ − *α*^−^) between the systole and diastole trials was computed as a measure of the cardiac cycle effect on learning rates.

## Results

### Manipulation check

Figure 2A illustrates the histogram detailing the presentation of discriminative stimuli in relation to the cardiac cycle. The precision of the onset timing in the cardiac cycle relative to the R-wave peak indicated that >99% of the trials were within 200 ms of the manipulated timing in both systole (red) and diastole (blue) trials. Specifically, the mean time from the R-wave peak for the systole trials was 244 ms (SE = 27 ms), whereas the mean time from the R-wave peak for the diastole trials was 527 ms (SD = 14 ms). The precise timing within the cardiac cycle was comparable to that reported in previous studies (e.g., Azevedo et al., 2017, 2018). Thus, the manipulation check suggested that the onset of the display of the discriminative stimuli was successfully synchronized to coincide with either the participant's cardiac systole or diastole.

**Figure 2.**
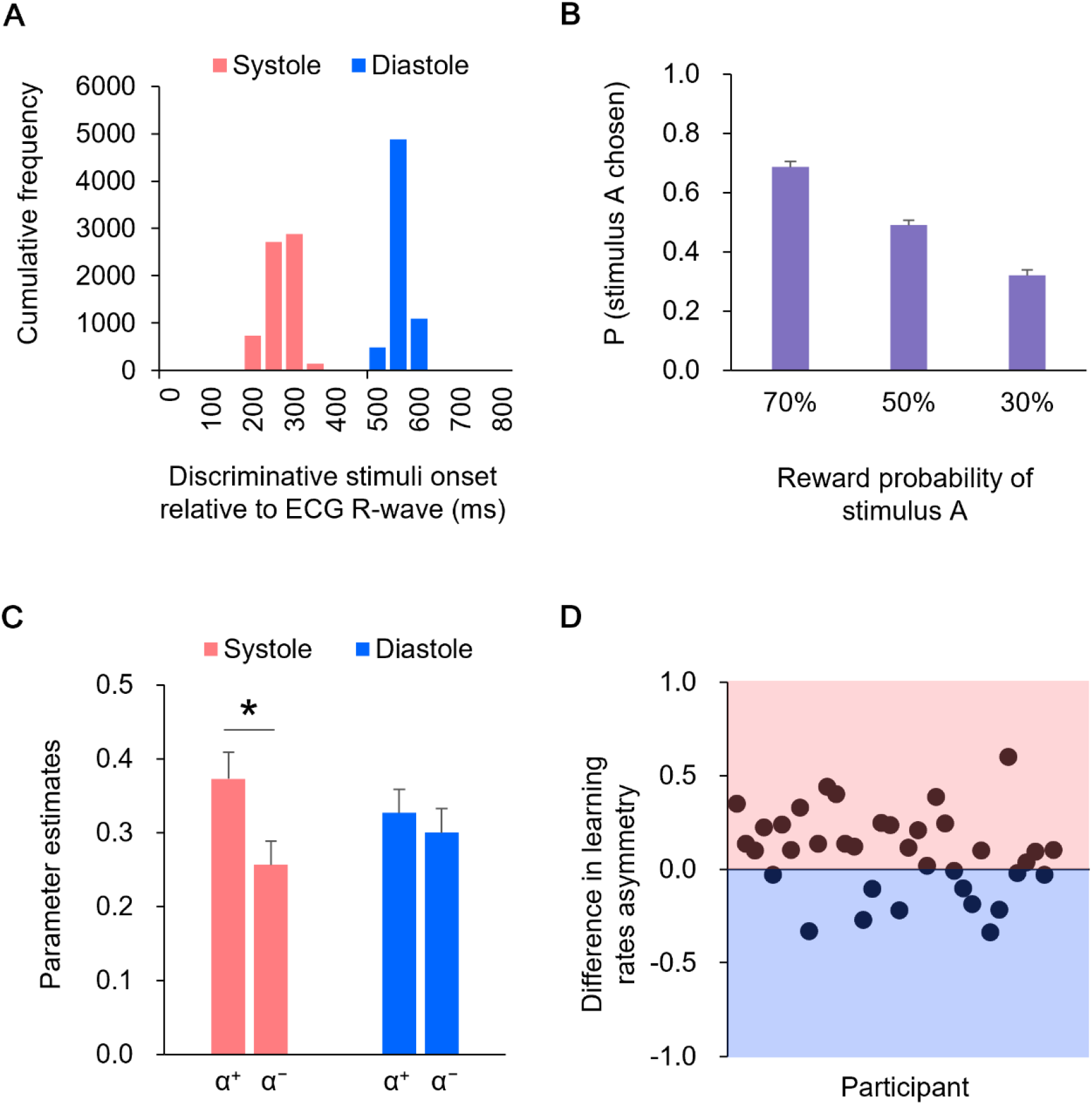
**(A)** The precision of the timing within the cardiac cycle, relative to the R-wave peak, is shown in the histogram. **(B)** The proportions of choosing stimulus A for each reward probability block. Error bars indicate *SE*. **(C)** Mean of the learning rates (*α*^+^ and *α*^−^) in the systole and diastole trials. Error bars indicate *SE*. An asterisk indicates a significant difference in the learning rate between the systole and diastole trials (*: *p* < .05). **(D)** The difference in the learning rate asymmetry (*α*^+^ − *α*^−^) between the systole and diastole trials for each participant. The positive value represents a larger learning rate asymmetry in the systole trials relative to the diastole trials, whereas the negative value represents a larger learning rate asymmetry in the diastole trials relative to the systole trials. In 24 of 36 participants, there were larger learning rate asymmetry in the systole trials compared to the diastole trials.

### Behavioral data

The mean reaction time was 624 ms (SE = 41 ms) in the systole trials and 741 ms (SE = 67 ms) in the diastole trials. A paired *t*-test revealed that the mean reaction time was shorter in systole trials than in diastole trials (*t*(35) = 2.42, *p* = .05, *d* = 0.40).

Figure 2B shows the proportions of choosing stimulus A for each reward probability block. As shown in Figure 2B, as the level of reward probability of stimulus A decreased, the proportions of choosing stimulus A decreased. One-way ANOVA (three reward probabilities) on the proportions of choosing stimulus A revealed a significant effect of reward probability (*F*(2,70) = 102.34, *p* < 0.01, partial η^2^ = 0.75). Post-hoc comparisons indicated that the differences in the proportions of choosing stimulus A were significant across all reward probabilities (*p*s < .01).

### Model-based analysis

Table 1 lists the log-marginal likelihoods of each model. We compared the log-marginal likelihood of each model and found that the Q-AF model had the highest value. Given that a marginal likelihood penalizes a complex model with extra parameters, and a higher marginal likelihood represents a better model, the Q-AF model was the best-fitting model for participants’ choice data in this study. We then estimated the parameters in the modified version of the Q-AF model in which the parameters of interest (*α*^+^, *α*^−^, and *β*) were allowed to have different values in the systole and diastole trials.

Figure 2C shows the learning rates (*α*^+^ and *α*^−^) in the systole and diastole trials. The learning rates were subjected to two-way repeated-measures ANOVA, with two trial types (systole and diastole) × two learning rate types ( + and −). The results revealed a significant interaction between trial and learning rate types (*F*(1,35) = 5.97, *p* < 0.05, partial η^2^ = 0.15). Post-hoc comparisons revealed that *α*^+^ was significantly higher than *α*^−^ in the systole trials (*t*(35) = 2.74, *p* < .01, *d* = 0.46), whereas the difference between *α*^+^ and *α*^−^ was not significant in the diastole trials (*t*(35) = 0.69, *p* = .50, *d* = 0.12). In addition, *α*^+^ did not differ between the systole and diastole trials (*t*(35) = 1.63, *p* = .11, *d* = 0.27), whereas *α*^−^ tended to differ between the systole and diastole trials (*t*(35) = 1.95, *p* = .06, *d* = 0.32). Figure 2D illustrates the difference in learning rate asymmetry ( *α*^+^ − *α*^−^) between the systole and diastole trials for each participant. The figure indicates that a positive value represents a larger learning rate asymmetry in the systole trials relative to the diastole trials, whereas a negative value represents a larger learning rate asymmetry in the diastole trials relative to the systole trials. As shown in Figure 2D, a larger learning rate asymmetry in the systole trials compared to the diastole trials was present in 24 of 36 participants. A paired *t*-test of the inverse temperature *β* revealed no significant difference between the systole and diastole trials (*t*(35) = 0.48, *p* = .63, *d* = 0.08).

## Discussion

This study aimed to investigate whether the cardiac cycle affects instrumental reward learning. To this end, we manipulated the onset of the presentation of discriminative stimuli such that they coincided with either cardiac systole or diastole across trials. The precision of the onset timing in the cardiac cycle showed that discriminative stimuli were successfully displayed at either cardiac systole or diastole (Figure 2A). The behavioral results showed that as the level of the reward probability of stimulus A decreased, the proportions of choosing stimulus A decreased (see Figure 2B), indicating that the participants performed the instrumental learning task with the goal of maximizing their monetary rewards. These results confirm the validity of cardiac cycle manipulation and our experimental task.

The main finding of this study was that the learning rate for positive prediction errors was higher than that for negative prediction errors in the systole trials, whereas learning rates did not differ between positive and negative prediction errors in the diastole trials. In this study, we timed the presentation of discriminative stimuli with the systolic (~300 ms after the R-peak) or diastolic (~550 ms after the R-peak) phases of the cardiac cycle. It has been suggested that the arterial baroreceptor signal is processed in the brain approximately 300 ms after the R peak (e.g., Edwards et al., 2007; Gray et al., 2009). Therefore, it would be reasonable to consider that the different patterns of the learning rates between the systole and diastole trials could be caused by the effects of cardiac afferent signals on the processing of discriminative stimuli in the brain. This can be consistent with the study of Waselius et al. (2018) demonstrating that the cardiac cycle modulated neural processing of the conditioned stimulus and influenced Pavlovian learning. From this point of view, this study extends previous research by showing that cardiac cycle affected not only Pavlovian learning but also instrumental reward learning.

Previous research on the cardiac cycle effect has accumulated evidence indicating that the processing of motivational/affective stimuli can be facilitated during cardiac systole (for a review, see Garfinkel & Critchley, 2016). Although the discriminative stimuli used in this study were inherently neutral, it is natural to assume that they acquired motivational/affective significance through instrumental learning. Previous studies have shown that cardiac afferent signals enhance the awareness (Garfinkel et al., 2014) and attentional processing (Azevedo et al., 2018) of motivational/affective stimuli. Given that both awareness (e.g., Manns et al., 2000; Skora et al., 2021) and attentional processing of discriminative stimuli (for a review, see Mackintosh, 1975) are closely related to value updating in learning, the present results can be interpreted as indicating that cardiac afferent signals facilitated the awareness and attentional processing of discriminative stimuli, which affected subsequent value updating in instrumental reward learning.

The learning rate for positive prediction errors was higher than that for negative prediction errors in systole trials, whereas learning rates did not differ between positive and negative prediction errors in diastole trials. This means that the effect of the cardiac cycle on learning was observed as a difference in learning rate asymmetry rather than as an overall difference in learning rates. The higher learning rate for positive prediction errors than for negative prediction errors in systole trials was consistent with the results of previous studies (e.g., Frank et al., 2004; Lefebvre et al., 2017). Specifically, Lefebvre et al. (2017) demonstrated that human choice behavior in an instrumental learning task can be captured by the RL model, implementing a higher learning rate for positive prediction errors than for negative prediction errors (i.e., optimistic learning rate asymmetry). The authors suggested that this learning rate asymmetry is involved in optimism bias: overestimation of the likelihood of positive events compared to that of negative events (e.g., Sharot, 2011). From this perspective, the present results indicate that cardiac afferent signals can enhance the expression of optimistic learning rate asymmetry. This seems consistent with previous findings that depressed individuals exhibit impaired cardiac interoceptive ability (for a review, see Eggart et al., 2019) and show an absence of optimism bias (e.g., Sharot, 2011). Future research should explore the association between the effect of the cardiac cycle on learning and cardiac interoceptive ability, for instance, using a heartbeat detection task (e.g., Kleckner et al., 2015).

Although this study cannot draw definitive conclusions regarding the neural mechanism underlying the effect of the cardiac cycle on instrumental reward learning, some plausible interpretations can be proposed. Converging evidence suggests that dopaminergic systems are involved in instrumental reward learning (for a review, see Niv, 2009; O’Doherty et al., 2015). In humans, previous studies using neuroimaging techniques have repeatedly reported that prediction errors in instrumental reward learning are associated with neural signals in the striatum, which is known to be the major dopaminergic target (e.g.,Niv et al., 2012; Schönberg et al., 2007). Furthermore, it has been reported that dopaminergic manipulations by the administration of dopamine agonists or antagonists could influence neural activity related to prediction error and learning in the reward learning paradigm (e.g., Pizzagalli et al., 2008; van der Schaaf et al., 2012), indicating the causal role of dopaminergic activity in instrumental reward learning. Previous studies have indicated that the phasic discharge of arterial baroreceptors during cardiac systole encodes the strength and timing of arterial pressure at each heartbeat, which is conveyed to brain areas such as the amygdala, anterior cingulate cortex, insular cortex, and striatum (for a review, see Critchley & Harrison, 2013). Importantly, Yang and Lin (1993) demonstrated that elevation of arterial baroreceptor signals can lead to an increase in striatal dopamine release. Therefore, our results suggest the possibility that the cardiac afferent signal modulates the neural responses to discriminative stimuli in the dopaminergic system, which influences value updating in instrumental reward learning. This possibility is supported by previous findings that endogenous fluctuations in dopaminergic activity during the presentation of decision options influence the propensity to take risks by enhancing phasic neural responses to decision options (Chew et al., 2019). To better understand the neural mechanisms underlying the effect of the cardiac cycle on instrumental reward learning, future work is necessary to test this possibility by combining experimental paradigms assessing instrumental reward learning with neuroimaging techniques.

One important limitation of this study is the potential effect of the cardiac cycle on action-making and reward/punishment feedback processing. Since this study aimed to examine the effect of the cardiac cycle at the presentation of discriminative stimuli in the instrumental learning task, we manipulated the onset of the stimuli to be synchronized to coincide with the cardiac systole or diastole. This inevitably results in asynchronization of the cardiac timing of action making and the acceptance of feedback. Previous studies have shown that the cardiac cycle at action-making and delivery of outcome influences the experience of controlling one’s body to cause desired effects in the environment (i.e., the sense of agency) (Herman & Tsakiris, 2020). Furthermore, a previous study reported that action feedback processing reflected in event-related brain potentials was modulated by the cardiac cycle in a gambling task (Kimura, 2019). Therefore, it might be possible that the different cardiac timings at action-making and the acceptance of outcomes have affected learning. However, it should be emphasized that we manipulated only the onset of the discriminative stimuli to avoid varying the temporal relationship between the onset of the discriminative stimuli, action making, and the onset of the outcome, as the temporal relationship among them is a critical determinant of learning (e.g., Gallistel & Gibbon, 2000). If future research could develop a solution to manipulate the cardiac cycle while maintaining a stable temporal relationship, the effects of the cardiac cycle on learning would become more apparent.

## Conclusion

This study demonstrated that the learning rate asymmetry, which was estimated using a computational RL model, can be affected by the cardiac cycle. In particular, we showed that the expression of optimistic learning rate asymmetry was enhanced when discriminative stimuli were displayed during cardiac systole. Our results provide evidence that the natural fluctuation of cardiac afferent signals modulates the awareness and attentional processing of discriminative stimuli, which affect asymmetric value updating in instrumental reward learning.

## Acknowledgments

The authors would like to thank Moena Okibuchi and Tomoko Otomo (AIST, Japan) for their assistance in conducting the experiments. This work was supported by a Grant-in-Aid for Scientific Research to K. Kimura from the Japan Society for the Promotion of Science (18K12023).

## Author contributions

K. Kimura designed the behavioral paradigms. K. Kimura and N.K. conducted the experiments and collected the data. K. Kimura, A.T., and K. Katahira analyzed the data. K. Kimura wrote the paper. All authors read and approved the final manuscript.

